# *Prochlorococcus* exudate stimulates heterotrophic bacterial competition with rival phytoplankton for available nitrogen

**DOI:** 10.1101/2021.08.25.457542

**Authors:** Benjamin C. Calfee, Liz D. Glasgo, Erik R. Zinser

**Affiliations:** Department of Microbiology, University of Tennessee, Knoxville, TN, USA

**Keywords:** *Prochlorococcus*, *Synechococcus*, *Alteromonas*, competition, nitrogen limitation

## Abstract

The marine cyanobacterium *Prochlorococcus* numerically dominates the phytoplankton community of the nutrient-limited open ocean, establishing itself as the most abundant photosynthetic organism on Earth. This ecological success has been attributed to lower cell quotas for limiting nutrients, superior resource acquisition, and other advantages associated with cell size reduction and genome streamlining. In this study we tested the prediction that *Prochlorococcus* outcompetes its rivals for scarce nutrients, and that this advantage leads to its numerical success in nutrient-limited waters. Strains of *Prochlorococcus* and its sister genus *Synechococcus* grew well in both mono- and co-culture when nutrients were replete. However, in nitrogen-limited medium *Prochlorococcus* outgrew *Synechococcus*, but only when heterotrophic bacteria were also present. In the nitrogen-limited medium, the heterotroph *Alteromonas macleodii* outcompeted *Synechococcus* for nitrogen, but only if stimulated by exudate released by *Prochlorococcus*, or if a proxy organic carbon source was provided. Analysis of a nitrate reductase mutant *Alteromonas* suggested that *Alteromonas* outcompetes *Synechococcus* for nitrate, during which co-cultured *Prochlorococcus* grows on ammonia or other available nitrogen species. We propose that *Prochlorococcus* can stimulate antagonism between heterotrophic bacteria and potential phytoplankton competitors through a metabolic cross-feeding interaction, and this stimulation could contribute to the numerical success of *Prochlorococcus* in the nutrient-limited regions of the ocean.

**Significance Statement:** In nutrient-poor habitats, the competition for limited resources is thought to select for organisms with enhanced ability to scavenge nutrients and utilize them efficiently. Such adaptations characterize the cyanobacterium *Prochlorococcus*, the most abundant photosynthetic organism in the nutrient-limited open ocean. In this study the competitive superiority of *Prochlorococcus* over a rival cyanobacterium, *Synechococcus*, was captured in laboratory culture. Critically, this outcome was achieved only when key aspects of the open ocean were simulated: a limited supply of nitrogen, and the presence of heterotrophic bacteria. Results indicate that *Prochlorococcus* promotes its numerical dominance over *Synechococcus* by energizing the heterotroph’s ability to outcompete *Synechococcus* for available nitrogen. This study demonstrates how interactions between trophic groups can influence interactions within trophic groups.

## Introduction

The phytoplankton community occupying the vast majority of the sunlit ocean experiences chronic nutrient limitation (1–4). Depending on the location, the limiting nutrient(s) include nitrogen, phosphorus, iron, and other metals. While the diversity of phytoplankton in these regions can be quite high, numerical superiority is often achieved by a single genus of cyanobacteria, *Prochlorococcus*. The most abundant photosynthetic organism in the ocean, *Prochlorococcus* can grow to populations that exceed 100,000 cells ml^-1^, besting their competitors by orders of magnitude in many instances (5–8).

The reasons underpinning the numerical dominance of *Prochlorococcus* in nutrient-limited waters have not been fully elucidated, but several distinguishing features of this unusual cyanobacterium have been implicated. *Prochlorococcus* has the smallest cell and genome size for a photoautotroph, which collectively lowers the cell quota for nitrogen, iron, and phosphorus (9–12). Phosphorus quota is further reduced by the replacement of phospholipids with sulfolipids as the predominant membrane lipids (13, 14). Additional means of economy (10, 15–17) may further contribute to the ability of *Prochlorococcus* to reproduce at lower cost relative to its competitors under nutrient-limited conditions.

Reduction in cell size is thought to provide *Prochlorococcus* with the additional advantage of superior nutrient acquisition (18). Lomas et al. 2014 noted that when normalized to cell quota, *Prochlorococcus* had a higher affinity to phosphorus relative to *Synechococcus* and picoeukaryotic phytoplankton (19). Notably, resource competition theory applied to global ocean simulations predicted the numerical domination of the oligotrophic regions by analogs of *Prochlorococcus*, which could draw nutrients down to concentrations that cannot be accessed by their competitors (20–22).

Despite the net loss of genes through streamlining, the diversity within *Prochlorococcus* is high and believed to contribute to the numerical dominance of *Prochlorococcus* by facilitating niche expansion. Phylogenetically distinct clades, termed ecotypes, exist within the genus and have demonstrated different optima for temperature, light intensities, and nutrient utilization that correlate with their environmental distributions (23–31). Notably, within these ecotypes, sub-ecotypes have been found with their own distinct ecologies, suggesting that the open ocean niche is finely partitioned through environmental influences on *Prochlorococcus* evolution (32–34).

A final contributor to the ecological success of *Prochlorococcus* may be the help it receives from the microbial community. All known genomes of *Prochlorococcus* lack the gene encoding the hydrogen peroxide scavenger, catalase (35–37). Loss of catalase is believed to improve growth efficiency by reducing cell quotas for iron and/or nitrogen, but it leaves cells highly susceptible to oxidative damage from environmental sources of hydrogen peroxide (12, 36, 38). *Prochlorococcus* survives this threat because it is cross-protected by co-occurring catalase-positive “helpers” such as *Alteromonas macleodii*, a heterotroph frequently co-isolated with *Prochlorococcus* (12, 35, 39). *Alteromonas macleodii* rapidly scavenges extracellular H_2_O_2_, causing changes in gene expression and promoting the growth of co-cultured *Prochlorococcus* in conditions that would otherwise be lethal (35, 40–42).

The physiological and genetic evidence all predict the success of *Prochlorococcus* over its competitors in the nutrient-limited ocean. In this work we sought direct evidence that *Prochlorococcus* could achieve numerical success over a key rival, *Synechococcus*. We focused our study on nitrogen-limiting conditions simulating the North Pacific Subtropical Gyre (NPSG) (43) where *Prochlorococcus* outnumbers *Synechococcus* and other rival phytoplankton by an order or magnitude or more (6, 8, 44). We found that competition for nitrogen explained the differences in *Prochlorococcus* and *Synechococcus* abundance, but only through the presence and specific activity of marine heterotrophic bacteria fed by *Prochlorococcus-derived* carbon. As these outcomes matched prior predictions of *Prochlorococcus* success, we argue that conditions such as the ones examined could provide important insight into the global ecology of *Prochlorococcus*.

## Results

### *Prochlorococcus* outcompetes *Synechococcus* in the presence of heterotrophs

Cyanobacterial growth in mono- and co-cultures was assessed in low-nitrogen medium (AMP-MN), an artificial seawater medium lacking N amendment, and containing approximately 0.4 μM residual bioavailable N (Fig. S1). *Prochlorococcus* strain MIT9215 reached a higher maximum abundance in monoculture than when in coculture with *Synechococcus* strain WH7803, suggesting that competition in coculture caused a slight but significant reduction in MIT9215 cell yield (Fig. 1A) (p <0.0001). Inversely, WH7803 maximum abundance was lower in monoculture than when in coculture with MIT9215, however this difference was not significant (Fig. 1A) (p = 0.2754).

**Figure 1.**
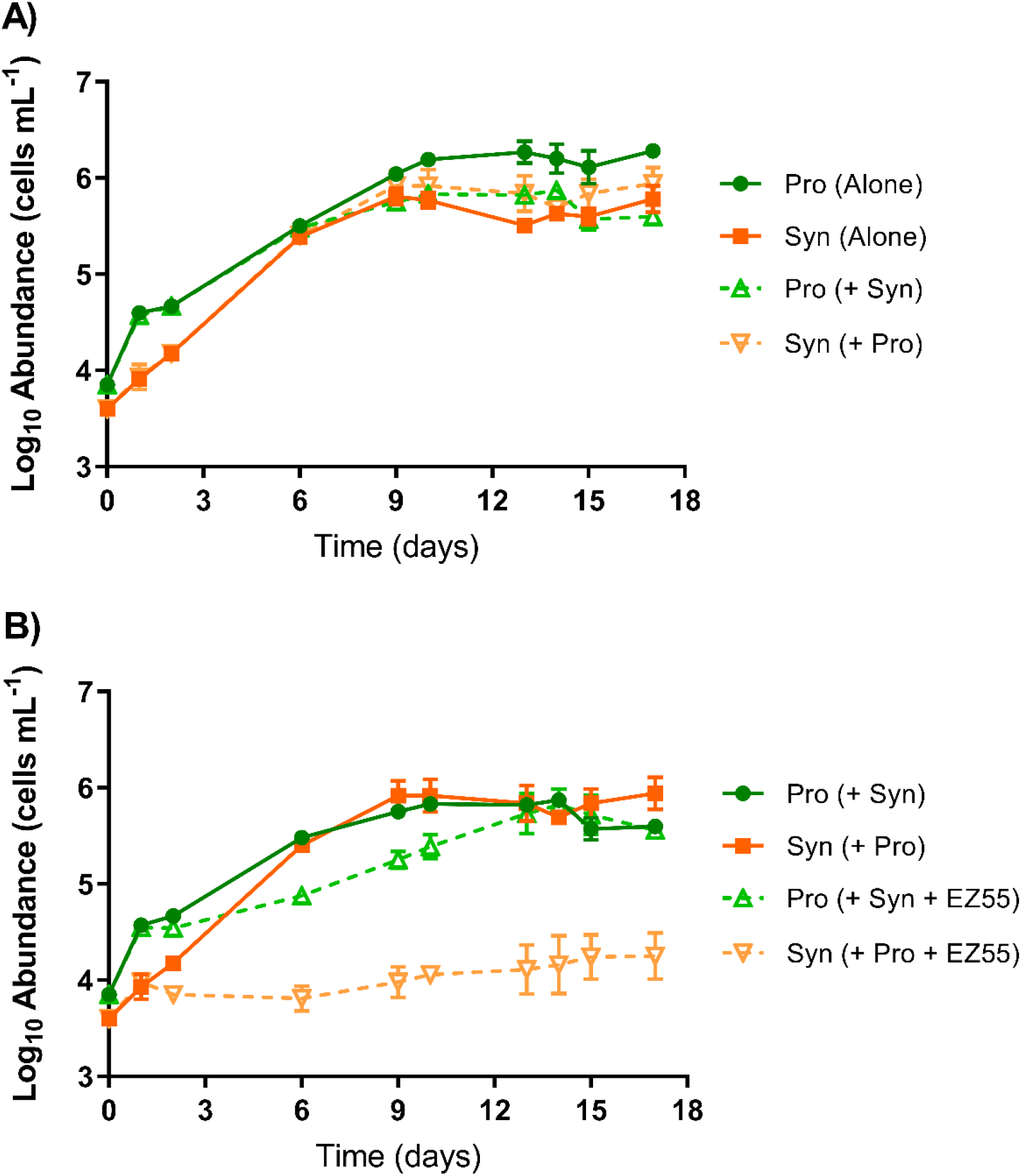
Growth of *Prochlorococcus* strain MIT9215 and *Synechococcus* strain WH7803 in AMP-MN artificial seawater medium in monoculture (A), cyanobacteria coculture (A and B), and tripartite culture with *Alteromonas macleodii* strain EZ55 (B). Error bars represent one standard deviation of the geometric mean (n=3).

Addition of the marine heterotrophic bacterium, *Alteromonas macleodii* strain EZ55, dramatically changed the outcome for the *Synechococcus*-*Prochlorococcus* cocultures (Fig. 1B). While MIT9215 growth rate declined moderately, addition of EZ55 to the coculture resulted in a near total loss of growth for WH7803 (p = 0.0018). In this AMP-MN medium, the EZ55 heterotroph grew rapidly to ~10^6^ cells mL^-1^, regardless of whether cyanobacteria were present (data shown in Fig. 3 and 4), indicating growth on trace contaminating organic carbon in the medium. The presence of the heterotroph in this nitrogen-limited medium thus shifted the phytoplankton community structure to one resembling open ocean communities, with *Prochlorococcus* numerically-dominant over rival *Synechococcus*.

The dynamics of resource competition were further investigated by challenging the cyanobacterial strains to invade established populations of their competitors when rare. At day 32 of growth in AMP-MN, a small inoculum (~3.00e^3^ cell mL^-1^) from WH7803 monocultures was added to cultures of MIT9215 ± EZ55; reciprocally, MIT9215 monocultures were inoculated into WH7803 ± EZ55 cultures. WH7803 were able to invade MIT9215 monocultures after a few days lag and reach an almost equal abundance over the next 17 days (Fig. 2A). However, WH7803 failed to grow in MIT9215 cultures when EZ55 was present, dropping below the limit of detection shortly after inoculation (Fig. 2B).

**Figure 2.**
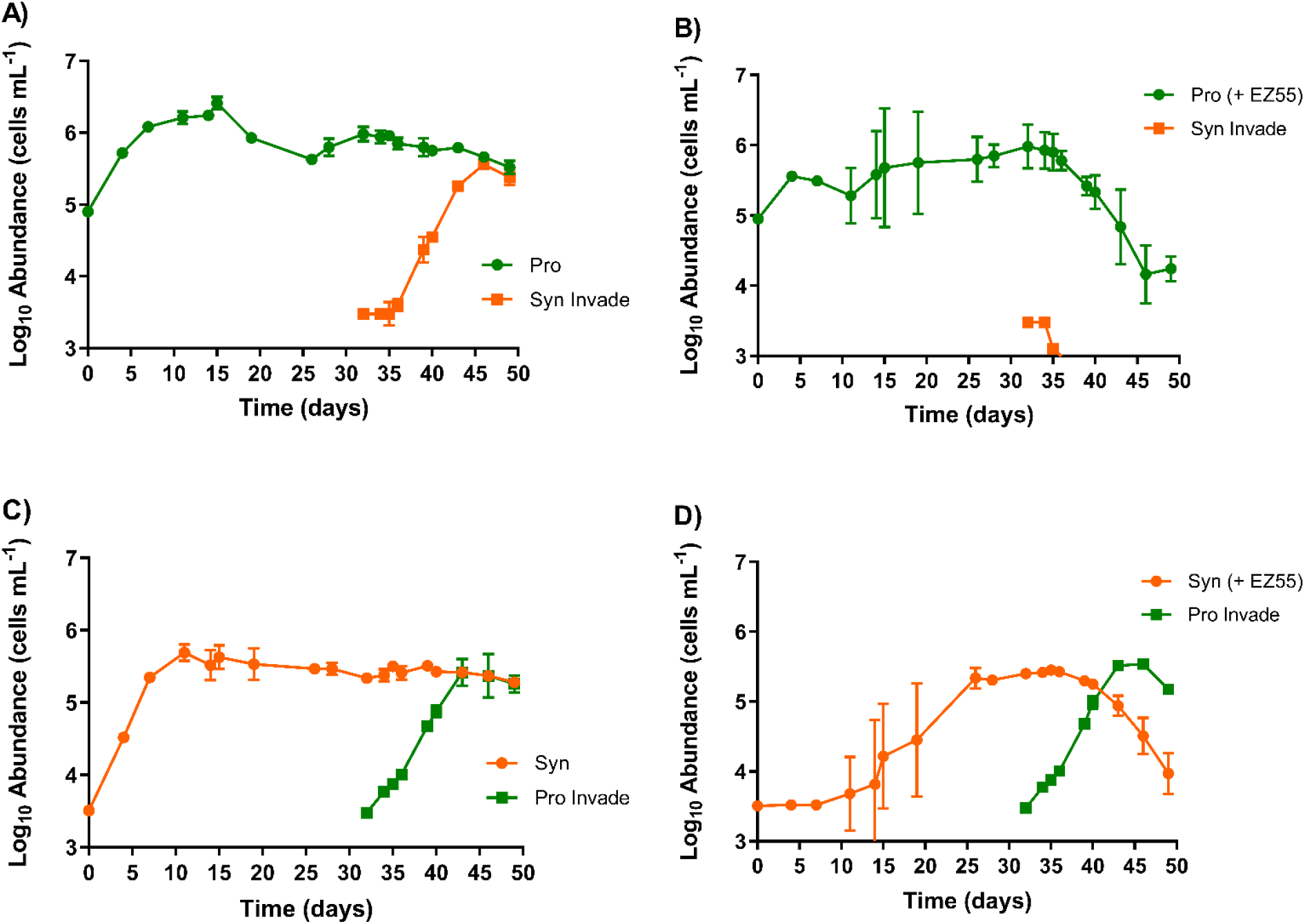
Growth of *Prochlorococcus* strain MIT9215 (A-B) and *Synechococcus* strain WH7803 (C-D) in AMP-MN artificial seawater medium with and without *Alteromonas macleodii* strain EZ55. On day 32, cultures of the cyanobacteria without *Alteromonas* were inoculated as minority into the cultures of the rival cyanobacterium with and without *Alteromonas* to assess ability to invade. Error bars represent one standard deviation of the geometric mean (n=3).

In the reciprocal invasion assay, MIT9215 rapidly grew when inoculated in the WH7803 monoculture, with both organisms co-existing at equal abundances (Fig. 2C). In the presence of EZ55, MIT9215 was still able to invade a culture of WH7803 (Fig. 2D). Interestingly, with EZ55 present, the MIT9215 population displaced WH7803 as the majority phytoplankter in the culture: WH7803 exhibited a dramatic decline in abundance (Fig. 2D) that was not observed when EZ55 was absent (Fig. 2C). Thus, independent of the starting ratios or cell concentration, the presence of the EZ55 heterotroph favored the growth of *Prochlorococcus* over *Synechococcus* when nitrogen was scarce.

### *Prochlorococcus* exudate drives heterotroph N competition with *Synechococcus*

Critically, the inhibitory effect of EZ55 on WH7803 growth was absent if the *Prochlorococcus* MIT9215 strain was not included. WH7803 growth showed no significant difference in growth between mono- and coculture with EZ55 in AMP-MN (Fig. 3A) (p = 0.91).

**Figure 3.**
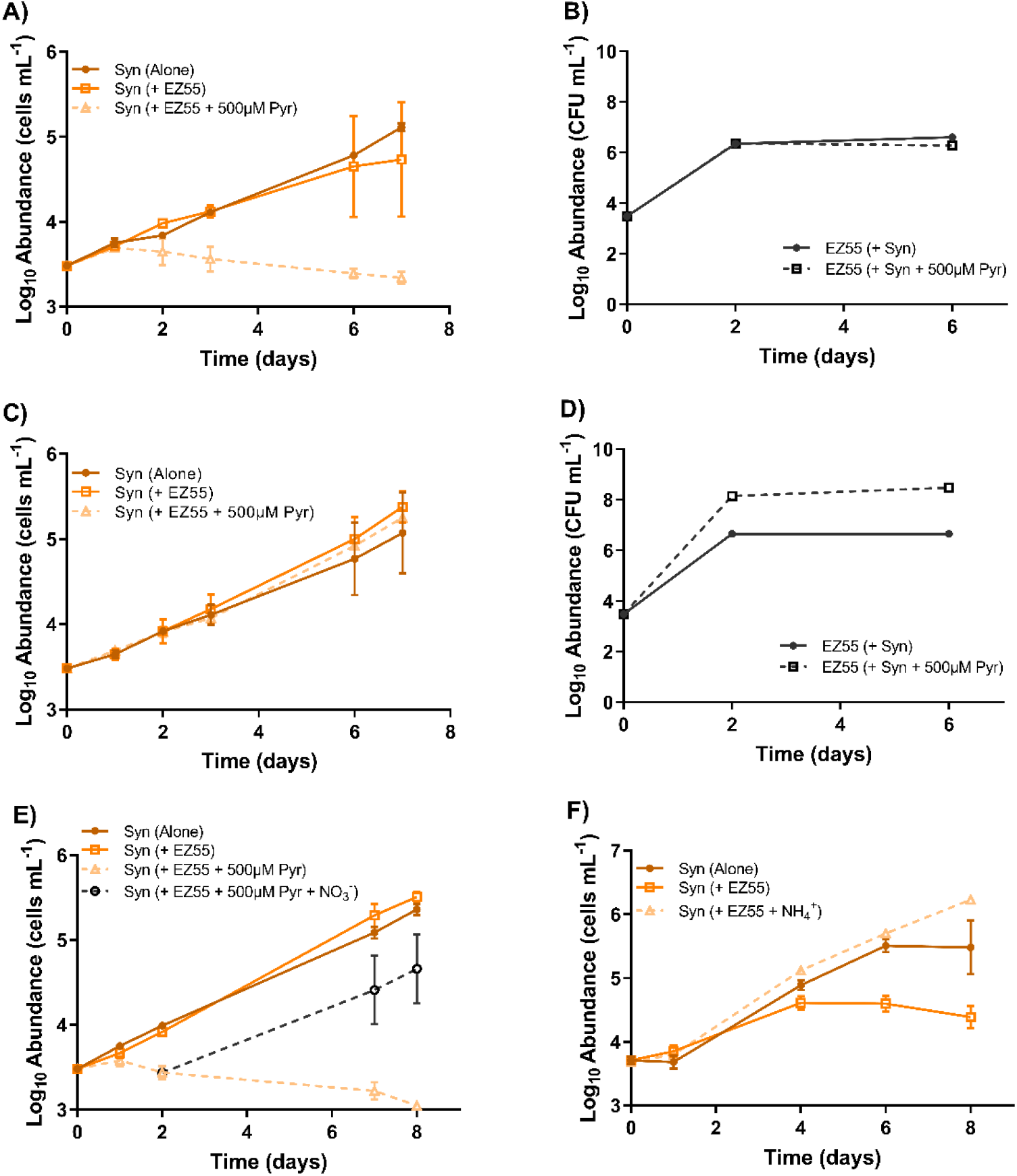
Growth of *Synechococcus* strain WH7803 (A, C, E, and F) and *Alteromonas macleodii* strain EZ55 (B and D) in AMP-MN (A, B, and E) and AMP-A (C and D) artificial seawater media in monoculture, coculture, and coculture with the addition of 500 μM sodium pyruvate (A-E). Cocultures were also amended with 500 μM sodium pyruvate and 800 μM sodium nitrate to demonstrate growth rescue by nutrient addition (E). WH7803 was also cultured in *Prochlorococcus*-conditioned AMP-MN alone and with EZ55 ± 400 μM NH_4_^+^ (F). Error bars represent one standard deviation of the geometric mean (n=3).

This outcome suggested that *Prochlorococcus* may be secreting a factor(s) that stimulates the competition of EZ55 for resource(s) shared by WH7803. To test this, EZ55 and WH7803 were placed in co-culture competition in media pre-conditioned by MIT9215. Whether MIT9215 remained (Fig. S2A and B) or were removed (via filtration) prior to competition (Fig. 3F), the outcome was the same: in contrast to unconditioned medium (Fig. 3A), WH7803 was inhibited in MIT9215-conditioned medium.

We next considered two hypotheses for the *Prochlorococcus*-driven loss of WH7803 growth in the presence of EZ55: either *Prochlorococcus* is driving EZ55 to compete for limited resources, or to produce a factor that is toxic to WH7803. Carbon and nitrogen amendment studies favored the former over the latter hypothesis.

*Prochlorococcus* releases a large fraction of fixed carbon as dissolved organic carbon during nitrogen-limited growth (45), so we reasoned that this excess source of carbon and energy could be energizing EZ55 to compete with *Synechococcus* for nitrogen in this nitrogen-limited medium. Pyruvate was examined as a proxy for *Prochlorococcus* exudate, and like the exudate, allowed EZ55 to prevent the growth of WH7803 (Fig. 3A). Notably, in tripartite cultures, addition of pyruvate (Fig. S3) further contributed to WH7803 reduction without apparent effect on MIT9215.

In the AMP-MN medium, nitrogen is the limiting resource for both *Prochlorococcus* and *Synechococcus* (Fig. S1A and S1B); other nutrients were provided in excess. As such, we reasoned that if EZ55 was restricting growth of WH7803, it was likely via competition for nitrogen. Consistently, addition of excess nitrogen to the medium as either NH_4_^+^ or NO_3_^-^ restored the ability of WH7803 to grow in the presence of pyruvate or exudate-stimulated EZ55, whether at the onset of co-cultivation (Fig. S2A, 3C, and 3F) or after WH7803 has ceased growth for several days (Fig. 3E). Notably, in these co-culture studies, pyruvate additions enabled EZ55 to grow to several orders of magnitude higher when nitrogen was in excess (Fig. 3D) but not when limiting (Fig. 3B), suggesting the inhibition by EZ55 requires excess carbon relative to nitrogen.

### Nitrogen Competition in Three-Member Cocultures

While the concentration of total bioavailable N in AMP-MN has been established (Fig S1), the constituent N species are not known. We hypothesized that while *Prochlorococcus* strain consumes the NH_4_^+^, the *Synechococcus* and heterotroph strains compete for a residual N resource *Prochlorococcus* cannot utilize but the other two can, namely NO_3_^-^ (46). To test this hypothesis we generated a transposon insertion mutant of EZ55 that lacks the nitrate reductase enzyme encoded by *nirB*. The *nirB* mutant cannot utilize nitrate as a nitrogen source, and, unlike wild type (Fig. 4A), cannot prevent growth of WH7803 in tripartite cultures with MIT9215 (Fig. 4B). The *nirB* mutation did not impact growth of the *Alteromonas* strain (Fig. 4C and 4D), suggesting this mutation prevented nitrogen competition without impacting overall growth. The inability of the nitrate utilization-deficient strain of EZ55 to restrict the growth of WH7803 suggests that NO_3_^-^ was present in AMP-MN, and that wild type EZ55 is able to outcompete WH7803 for this resource (when activated by *Prochlorococcus* exudate).

**Figure 4.**
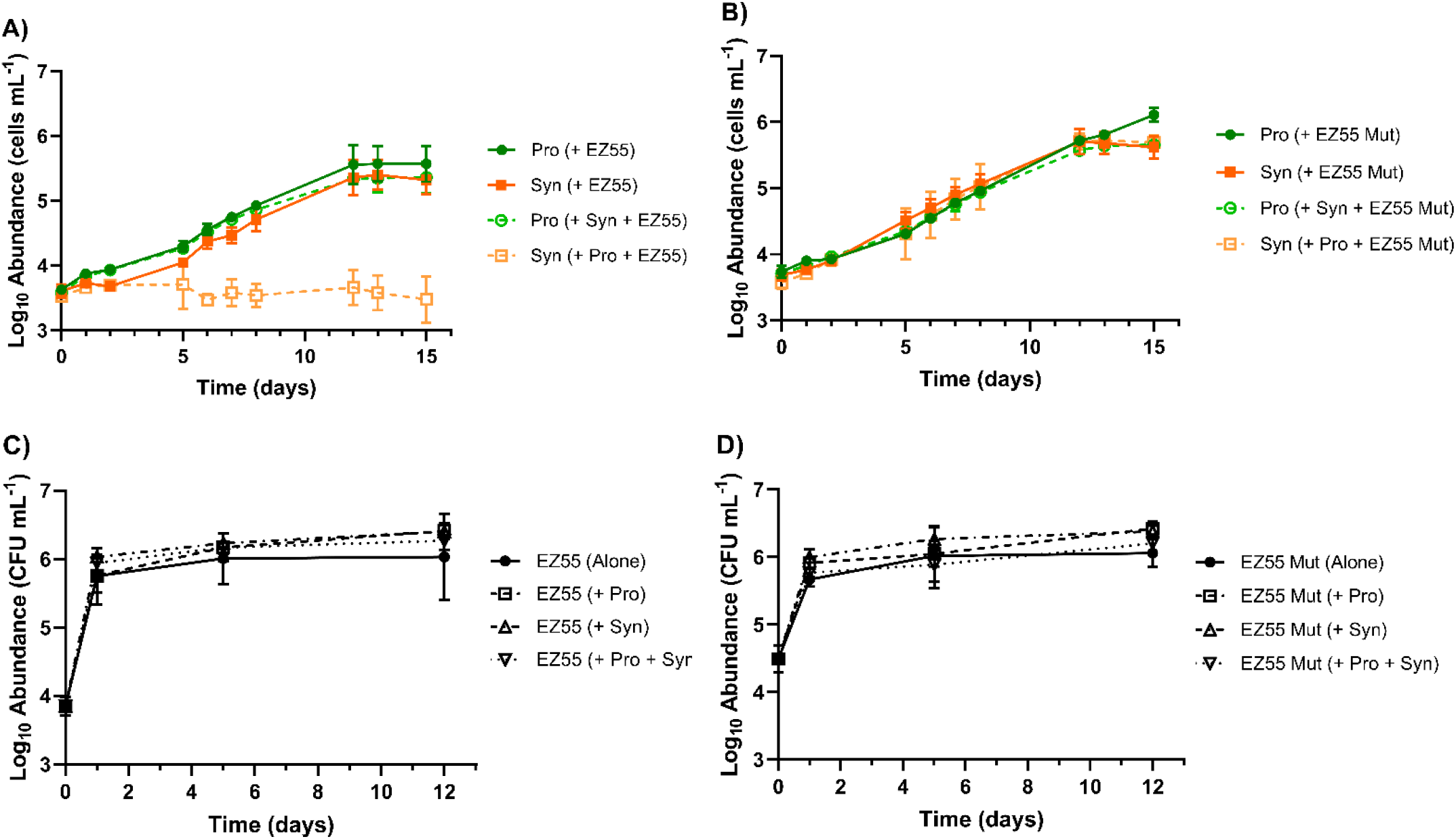
Growth of *Prochlorococcus* strain MIT9215 and *Synechococcus* strain WH7803 in AMP-MN artificial seawater medium in coculture and tripartite culture with *Alteromonas macleodii* strain EZ55 WT (A) or *Alteromonas macleodii* strain EZ55 mutant (B). Abundance of heterotroph in each treatment is shown for WT (C) and mutant (D). Error bars represent one standard deviation of the geometric mean (n=3).

### Competition outcomes are robust with regard to genotype

To determine the extent to which strain genotype impacts the outcomes of co-cultivation, we modified the mixed culture experiments by substituting MIT9215, WH7803, or EZ55 with different strains of *Prochlorococcus, Synechococcus*, or heterotrophic bacterium, respectively. Like MIT9215, high light-adapted *Prochlorococcus* strains MIT9312 or MED4 outcompeted WH7803 in the presence of EZ55 (Fig. 5A). And, like WH7803, *Synechococcus* strains CC9605 and WH8102 were outcompeted by MIT9215 in the presence of EZ55 (Fig. 5B).

**Figure 5.**
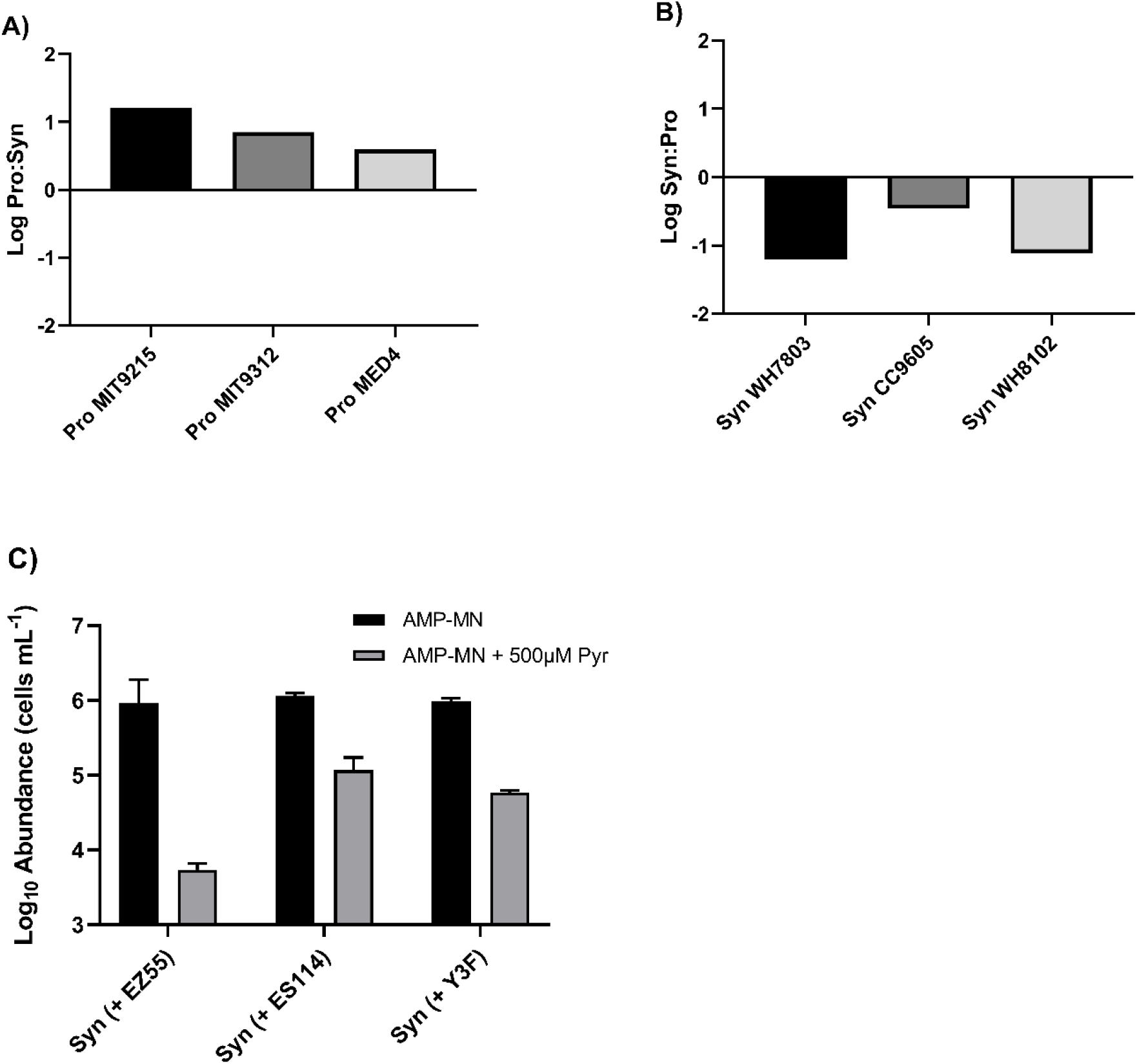
Log ratio of *Prochlorococcus* and *Synechococcus* strains maximal abundances in tripartite culture with *Alteromonas macleodii* strain EZ55 in AMP-MN artificial seawater medium. *Prochlorococcus* strains (A) were cultured with *Synechococcus* strain WH7803 and EZ55 and *Synechococcus* strains (B) were cultured with *Prochlorococcus* strain MIT9215 and EZ55. Maximum abundances of Synechococcus strain WH7803 were observed when cultured in AMP-MN or AMP-MN + 500 μM sodium pyruvate with different marine heterotrophic bacteria (C). Error bars represent one standard deviation of the geometric mean (n=3).

As a final constraint on the *Synechococcus* – heterotroph coculture outcomes, different marine heterotrophic bacteria were substituted for EZ55: *Phaeobacter* sp. strain Y3F and *Vibrio fischeri* strain ES114. When grown in N-replete AMP-A +/- pyruvate, or N-limited AMP-MN without pyruvate, coculturing with any of the three heterotrophs did not cause any significant deviation of WH7803 maximal abundance compared to the monoculture control (Fig. S4A, S4B, S4C). However, as with EZ55, the addition of pyruvate to AMP-MN caused a reduction in WH7803 maximal abundance when in co-culture with YF3 or ES114 compared to either the monoculture control (Fig. S4D) (p <0.0001) or cocultures in AMP-MN without pyruvate (Fig. 5C) (p <0.0001). With the exception of ES114, all heterotrophs maintained steady long-term populations in AMP-MN regardless of amendments; ES114 declined steadily and only maintained its starting abundance with pyruvate addition (Fig. S4E-S4G).

## Discussion

The numerical dominance of *Prochlorococcus* over rival phytoplankton in the oligotrophic ocean has been recognized since its discovery in 1988 (47). In this study we describe conditions in which such dominance is reproduced in culture. Importantly, we observed that *Prochlorococcus* outgrows *Synechococcus* in low-nitrogen conditions, simulating the North Pacific Subtropical Gyre, and only in the presence of heterotrophic bacteria, simulating the multi-trophic mixed community of the ocean.

In the NPSG, where nitrogen is thought to limit growth (3, 4, 13, 48), *Prochlorococcus* can outnumber *Synechococcus* (and other members of the phytoplankton community) by several orders of magnitude (6, 8, 44). In these nitrogen-limited waters heterotrophic bacteria can grow to between 300,000 and 500,000 cells ml^-1^ and outnumber phytoplankton (49–51). Our low-nitrogen culture medium recapitulated these trends: heterotrophs grew to an only slightly greater abundance of 10^6^ cells ml^-1^, and in tripartite cultures the dynamics of the picocyanobacteria favored *Prochlorococcus* over *Synechococcus*, regardless of the relative starting abundances. When co-inoculated with *Alteromonas, Prochlorococcus* could grow while *Synechococcus* could not. In absence of *Prochlorococcus, Synechococcus* could grow and stably coexist with *Alteromonas* for weeks. However, once introduced into this culture, *Prochlorococcus* could invade rapidly and ultimately displace the *Synechococcus* population.

Our results suggest that *Prochlorococcus* acts indirectly, through a heterotroph intermediate, to dictate the growth outcome of its rival *Synechococcus* in low-nitrogen environments. In low nitrogen, low organic carbon medium, *Prochlorococcus* scavenges a residual source(s) of nitrogen, apparently with superior capability relative to *Alteromonas* and *Synechococcus*. *Alteromonas* can grow on residual organic carbon until it becomes growth arrested by a lack of carbon and energy. In this state it is poised to compete for nitrogen but lacks the carbon and energy resources to do so, unless fed by *Prochlorococcus*. Once fed, *Alteromonas* can begin to compete with *Synechococcus* for alternative nitrogen source(s). The inability of a mutant *Alteromonas* lacking the nitrate reductase enzyme to arrest the growth of *Synechococcus* suggests that the competition involves nitrate, a resource both *Synechococcus* and wild type *Alteromonas* can utilize, but the strains of *Prochlorococcus* examined in this study cannot. Nitrate-utilizing strains of *Prochlorococcus* have been recently isolated (52), and future studies in tripartite cultures with these strains could prove informative. In the paragraphs that follow we unpack this model to discuss the key supporting evidence and identify unanswered questions.

Our study implicates the release of organic carbon by *Prochlorococcus* for the stimulation of *Alteromonas* to outcompete *Synechococcus* for nitrogen. Neither *Prochlorococcus* nor *Alteromonas* acting alone was sufficient to diminish the growth of *Synechococcus*, but when together in a tripartite community diminished *Synechococcus* growth.

Importantly this effect was observed only when nitrogen was limiting in the medium; addition of excess nitrogen was all that was needed to retore *Synechococcus* growth. This latter result also argues against the production of a growth-limiting substance by *Alteromonas* as the explanation of the growth arrest of *Synechococcus*.

*Prochlorococcus* exudate was sufficient to stimulate the N-competition by *Alteromonas*, as was a proxy form of *Prochlorococcus* exudate, pyruvate. *Prochlorococcus* exudes a large fraction of fixed carbon as dissolved organic matter (53–55) much of which is bioavailable to heterotrophic bacteria (56, 57). Recently it was observed that *Prochlorococcus* can also release membrane vesicles (58), which may serve as complex nutrients for co-occurring heterotrophs. Critically, under nitrogen limitation the release of dissolved organic matter by *Prochlorococcus* is exacerbated (45, 59). The specific form(s) of released organic carbon that stimulated *Alteromonas* competition for nitrogen in this study is not known, but it is rather curious that

*Synechococcus* exudate was not sufficient for this effect: bipartite cultures of *Alteromonas* and *Synechococcus* stably co-existed in low-N medium. *Synechococcus* is known to release organic carbon, and this release increases under nutrient limitation (60), so this distinction between *Prochlorococcus* and *Synechococcus* exudate warrants further investigation.

As with carbon, the nitrogen species involved in the tripartite interactions are not yet completely identified. Our artificial seawater medium lacked nitrogen amendment, but trace amounts of nitrogen from unknown sources were capable of supporting microbial growth to 10^6^ cells ml^-1^. Due to the volatility of ammonia and reported cases of ammonia contamination in other systems (61), we suspect that it serves as a major component of the unamended medium. As the preferred nitrogen source for *Prochlorococcus* and most microbes, we suspect that ammonia is the primary nitrogen source consumed by *Prochlorococcus*, whether in mono- or mixed-cultures. However, strain MIT9215 has the genetic potential to utilize urea as well (37, 46), so this species cannot be ruled out. Nitrate is a likely component of the medium, as the ability of *Alteromonas* to prevent *Synechococcus* growth was eliminated when the nitrate reductase of the heterotroph was knocked out. While some strains of *Prochlorococcus* can utilize nitrate (52), the ones assayed in this study could not. Whether or not the nitrate-utilizing *Prochlorococcus* strains can also compete with *Synechococcus* for this resource could be resolved in future studies.

In the ocean, *Prochlorococcus* and *Synechococcus* compete for a variety of nitrogen sources, including organic forms such as amino acids (29, 62–66). In a 2018 study, Berthelot et al. observed that co-occurring populations of *Prochlorococcus, Synechococcus*, and the photosynthetic picoeukaryotes in the N-limited North Pacific Subtropical Gyre all utilize ammonia, urea, and nitrate, although to different extents (63).

While capable of sourcing their nitrogen from organic carbon molecules like amino acids, marine heterotrophs have been shown to also compete with phytoplankton for inorganic nitrogen in the form of ammonia or nitrate (67–69). Heterotrophs can account for 30% or more of inorganic nitrogen uptake at some locations (70, 71), and in some studies, inorganic nitrogen accounted for half or more of the total nitrogen acquired by heterotrophs (72, 73).

Importantly, the ability of heterotrophs to compete for inorganic nitrogen scavenging appears to be stimulated by organic carbon. Several studies by the Kirchman group and others noted the necessity of sufficient carbon for inorganic N uptake by bacteria (68, 69, 73–76). The physiological basis for this stimulation is not yet understood, however, studies with laboratory cultures provide some clues. For *Escherichia coli*, carbon limitation depletes the TCA cycle intermediate and key substrate for inorganic nitrogen assimilation, α-ketoglutarate (2-oxo-glutarate) (77). Consequently, C-starved cells have diminished rates of ammonium assimilation and potentially other N utilization pathways, even when cells are N limited (77). Notably, a recent study found that *Alteromonas* significantly reduced expression of genes involved in nitrogen metabolic pathways under carbon and iron colimitation (78).

The stimulation of inorganic nitrogen uptake in these studies is entirely consistent with our observations of *Alteromonas* and other marine heterotrophs in the N-limited medium. Like *E. coli*, carbon-limited *Alteromonas* may be deprived of the necessary α-ketoglutarate for assimilation of ammonia or nitrate. Alternatively, or in addition, carbon limitation may deprive the cells of the energy needed to drive transport of these substrates. In either case, the provision of organic carbon by *Prochlorococcus* appears to satisfy the requirements for enhanced inorganic nitrogen uptake and assimilation by these heterotrophs, out-competing *Synechococcus* in the process.

Prior studies have highlighted the beneficial effects of heterotroph interactions with picocyanobacteria (40–42, 60, 79–82). Previously we described how heterotrophic bacteria protect *Prochlorococcus* from oxidative stress (12, 38). Coe et al. (83) and Roth-Rosenberg et al. (84) have shown that heterotrophs promote the survival of *Prochlorococcus* during long-term light and nutrient (N or P) deprivation, respectively. Christie-Oleza et al. (60) found a similar relation between *Synechococcus* and a marine roseobacter. In that study, long-term co-existence under nutrient limitation was facilitated by an exchange of resources between the phototroph and heterotroph.

Interactions between picocyanobacteria have been less well characterized, but a recent study from Knight and Morris (85) showed that *Synechococcus* could aid the growth of *Prochlorococcus* under conditions simulating ocean acidification. The mechanism of this help was not identified, but because these co-cultures were grown in the presence of *Alteromonas* sp. EZ55, the authors speculated that *Synechococcus* could help *Prochlorococcus* indirectly by stimulating EZ55. The potential for allelopathic interactions between picocyanobacteria has also been noted (86–88).

Our study provides a new dimension to the picocyanobacteria-heterotroph and picocyanobacteria-picocyanobacteria interactions: the ability of one phototroph (*Prochlorococcus*) to drive a shift from co-existence to competition between a second phototroph (*Synechococcus*) and a heterotroph. Christie-Oleza et al. (60) found that *Synechococcus* and heterotroph strains co-exist during prolonged coculture in unamended seawater, and that upon N addition, cross-feeding could occur by the conversion of N substrates unusable by the other microbe: the heterotroph strain could convert organic nitrogen (peptone) to ammonia, while WH7803 could convert nitrate to dissolved organic nitrogen. In our study, both heterotroph and phototroph could utilize nitrate, and unless the former was mutated in its nitrate reductase, the heterotroph could apparently outcompete the *Synechococcus* strain for this resource when fed organic carbon by *Prochlorococcus*.

## Conclusion

This study demonstrates that metabolic interactions between trophic groups can influence relative abundances within trophic groups. The prediction that *Prochlorococcus* outcompetes rival phytoplankton including *Synechococcus* under nutrient limitation is largely confirmed, but this outcome may require the ability of *Prochlorococcus* to energize heterotrophic bacteria to outcompete their photosynthetic rivals for resources that they themselves do not use. If our results can be extrapolated to the natural environment, it highlights an important connection between carbon and nitrogen availability, and suggests complex microbial interactions can benefit streamlined, efficient genera such as *Prochlorococcus* to the detriment of their competition.

## Methods

### Strains and Culturing

Axenic cultures of *Prochlorococcus* strains MIT9215, MIT9312, and MED4, and *Synechococcus* strains WH7803, CC9605, and WH8102 were used in this study. Stock cultures of cyanobacteria were initially maintained in an artificial seawater medium, AMP-A (12, 89, 90), and were inoculated and serially maintained (for up to two years) in AMP-MN (this study, described below) to prevent introduction of excess nitrogen (N). Axenicity of cyanobacterial stocks and experimental cultures was tested routinely by diluting a small volume of culture into 1/10X ProAC and YTSS media and incubating these cultures in the dark at room temperature for up to six weeks to monitor for any increase in turbidity indicating presence of heterotrophic bacteria (35). All experiments were carried out at 24 °C in Percival I36VLX incubators (Percival, Boone, IA) with modified controllers that allowed for gradual increase and decrease of cool white light to simulate sunrise and sunset with peak midday light intensity of 150 μmol quanta m^-2^s^-1^ on a 14 hr:10 hr light:dark cycle (91). Ammonium (NH_4_^+^) was the N amendment in all experiments, unless otherwise stated, as it can be used by all strains in this study. Experiments that included different NH_4_^+^ concentrations were performed with NH_4_^+^ amendments to the AMP-A derivative, AMP-MN (<u>M</u>inus <u>N</u>itrogen), which is identical to AMP-A except that no N source is included. Stepwise amendments of NH_4_^+^ to AMP-MN indicated that the residual N bioavailable to *Prochlorococcus* and *Synechococcus* was approximately 0.4 μM (Figure S1).

Axenic heterotrophic bacteria utilized were *Alteromonas macleodii* strain EZ55 (35), *Vibrio fischeri* strain ES114 (92), and *Phaeobacter* sp. strain Y3F (93). Overnight cultures of heterotrophs were inoculated from cryo-preserved stocks prior to each experiment (−80°C in YTSS + 10% glycerol) into 5 mL volumes of YTSS (94) incubated shaking at 140 RPM at 24°C. Before inoculation into cyanobacterial cultures, heterotrophs were washed three times in 1.5 mL microcentrifuge tubes by centrifugation at 8,000 RPM for two minutes in a tabletop microcentrifuge and resuspension in 1mL AMP-MN.

While all culture media was sterilized by autoclaving, sterilized spent or *Prochlorococcus*-conditioned media was generated by culturing *Prochlorococcus* strain MIT9215 in large volumes of AMP-MN (~300 mL). At stationary phase these cells were removed by gentle filtration (−7 inHg) in a 1 L filter tower (Nalgene) using 0.2 μm pore size GTTP Isopore Membrane Filters (MilliporeSigma, Burlington, MA). Prior studies indicated low pressure filtration does not cause detectable rupture of *Prochlorococcus* cells during filtration (12). Sterility of this conditioned media was determined by flow cytometry alongside the experiments it was utilized in, in addition to the purity assay detailed above.

### Quantification of Cyanobacteria and Heterotroph Abundances

Abundances of cyanobacteria were quantified by flow cytometry using a Guava EasyCyte 8HT flow cytometer (Millipore, Burlington, MA) with populations of *Prochlorococcus* and *Synechococcus* differentiated in co-cultures by their red and red / yellow fluorescence, respectively (35, 95). Heterotrophs in mono and coculture experiments were quantified by viable counting with serial dilutions on YTSS 1.5% agar plates incubated at 24 °C.

### Transposon Mutagenesis

Mutants of *Alteromonas macleodii* strain EZ55 incapable of growing on nitrate (NO_3_^-^) as a sole N source were generated by transposon mutagenesis using a mini-Himar1 *Mariner* transposon carrying a kanamycin resistance selectable marker (96). The RB1 plasmid vector containing the transposon was propagated in *Escherichia coli* strain WM3064, a pir^+^ and 2,6-diaminopimelic acid (DAP) auxotroph donor strain (97). Overnight cultures of donor strain were inoculated from cryopreserved stocks (−80°C in LB + 10% glycerol) into 5 mL LB amended with 10 μg/mL kanamycin and 150 μL of 100 mM DAP (Alfa Aesar, Haverhill, MA) and incubated shaking at 37 °C. Conjugations with EZ55 were performed by plating both donor and recipient onto YTSS agar plates for 8 hours. Ex-conjugants were selected on YTSS + 10 μg/mL kanamycin agar plates. Selected colonies were screened for NO_3_^-^ utilization by replica plating (98) on AMP-A agar with 1.5% Noble agar (Difco) amended with 500 μM sodium pyruvate (Sigma-Aldrich) and either 400μM NH_4_^+^ or 882μM NO_3_^-^ as the nitrogen source. Replica plated colonies growing solely on plates containing NH_4_^+^ were transferred again into tubes of AMP-A with excess carbon and either nitrogen source to confirm the mutants were unable to grow on nitrate. Insertion location of the Mariner transposon within the nirB gene was verified by Arbitrary PCR (99), Sanger sequencing, and BLAST comparisons with the EZ55 genome (accession number 2785510739).

## Acknowledgements

We thank David Talmy and Elizabeth Fozo for helpful comments on the manuscript. We thank Jeff Gralnick and Jeff Morris for providing the pRB1 plasmid and WM3064 strains, respectively. This work was supported by NSF grants IOS-1451528 and OCE-2023680 to E.R.Z.

