## Supplemental Information for "*Prochlorococcus* exudate stimulates heterotrophic bacterial competition with rival phytoplankton for available nitrogen"

**This PDF file includes:**

Figures S1 to S4

**Supplemental Figures**

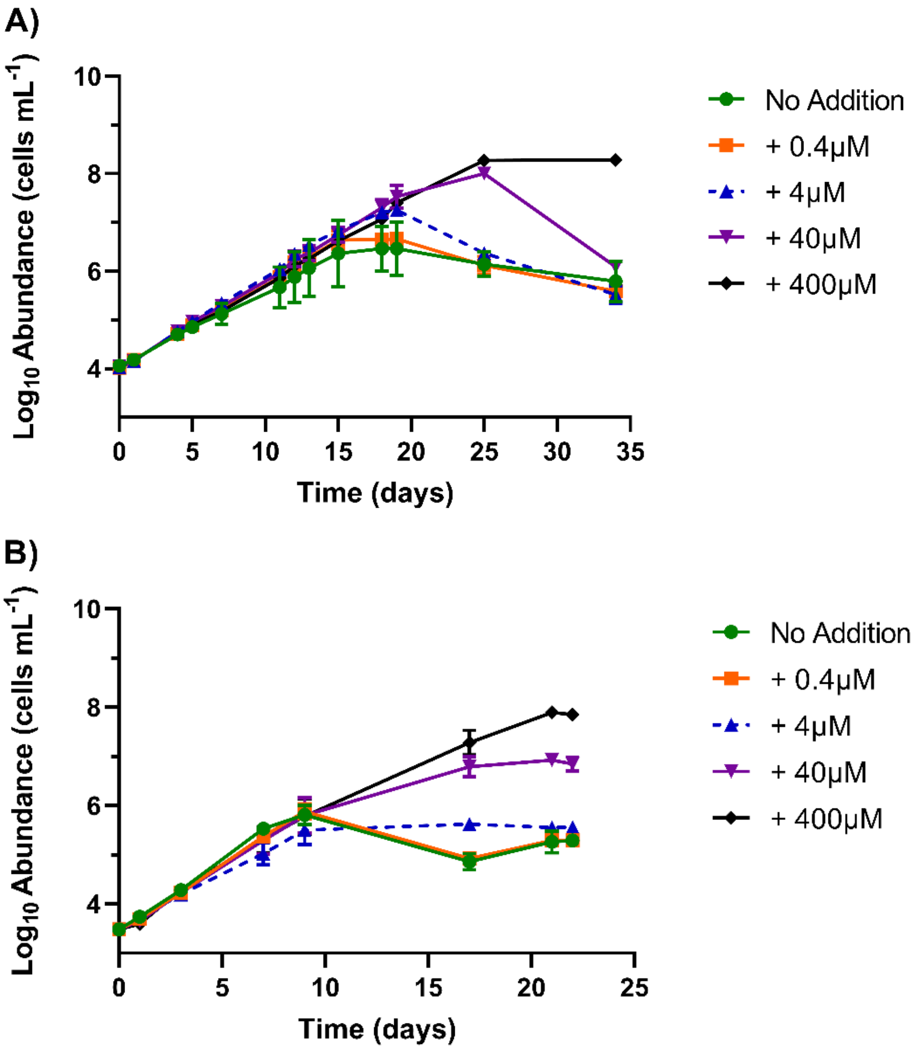


**Fig. S1** – Growth of *Prochlorococcus* strain MIT9215 (A) and *Synechococcus* strain WH7803 (B) in AMP-MN artificial seawater medium amended with varying concentrations of ammonium. Error bars represent one standard deviation of the geometric mean (n=3).

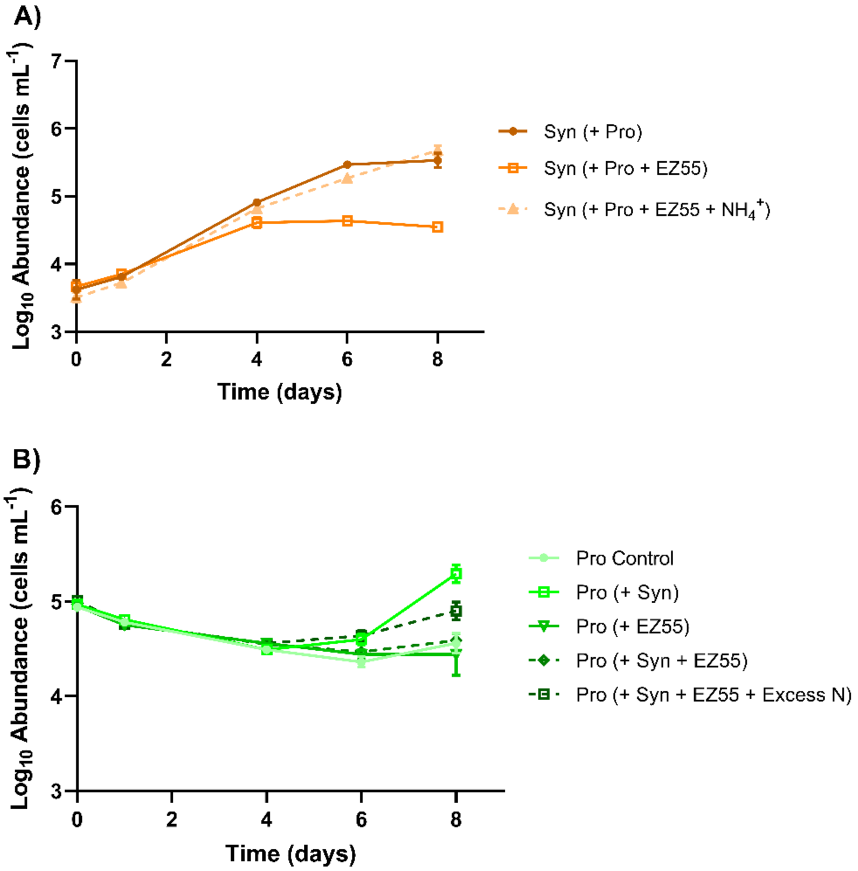


**Fig. S2** – Growth of *Synechococcus* strain WH7803 in AMP-MN artificial seawater medium pre-conditioned by *Prochlorococcus* strain MIT9215, which was still present during experiment. *Synechococcus* was inoculated alone and with *Alteromonas macleodii* strain EZ55 ± 400 µM $NH_4^+$ (A). Growth of *Prochlorococcus* strain MIT9215 in pre-conditioned AMP-MN artificial medium after inoculation of WH7803 and EZ55 (B). Error bars represent one standard deviation of the geometric mean (n=3).

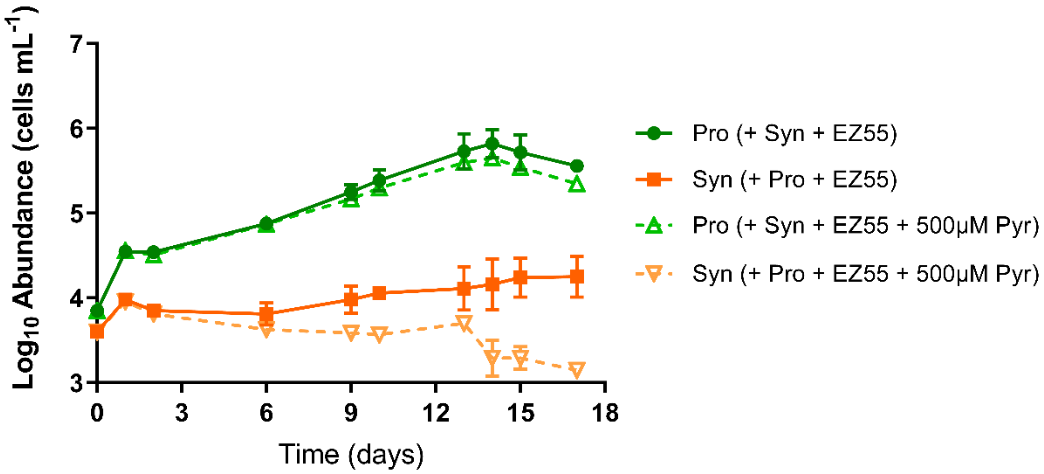


**Fig. S3** – Growth of with *Prochlorococcus* strain MIT9215, *Synechococcus* strain WH7803, and *Alteromonas macleodii* strain EZ55 in tripartite culture in AMP-MN artificial seawater medium with and without the addition of 500μM sodium pyruvate. Error bars represent one standard deviation of the geometric mean (n=3).

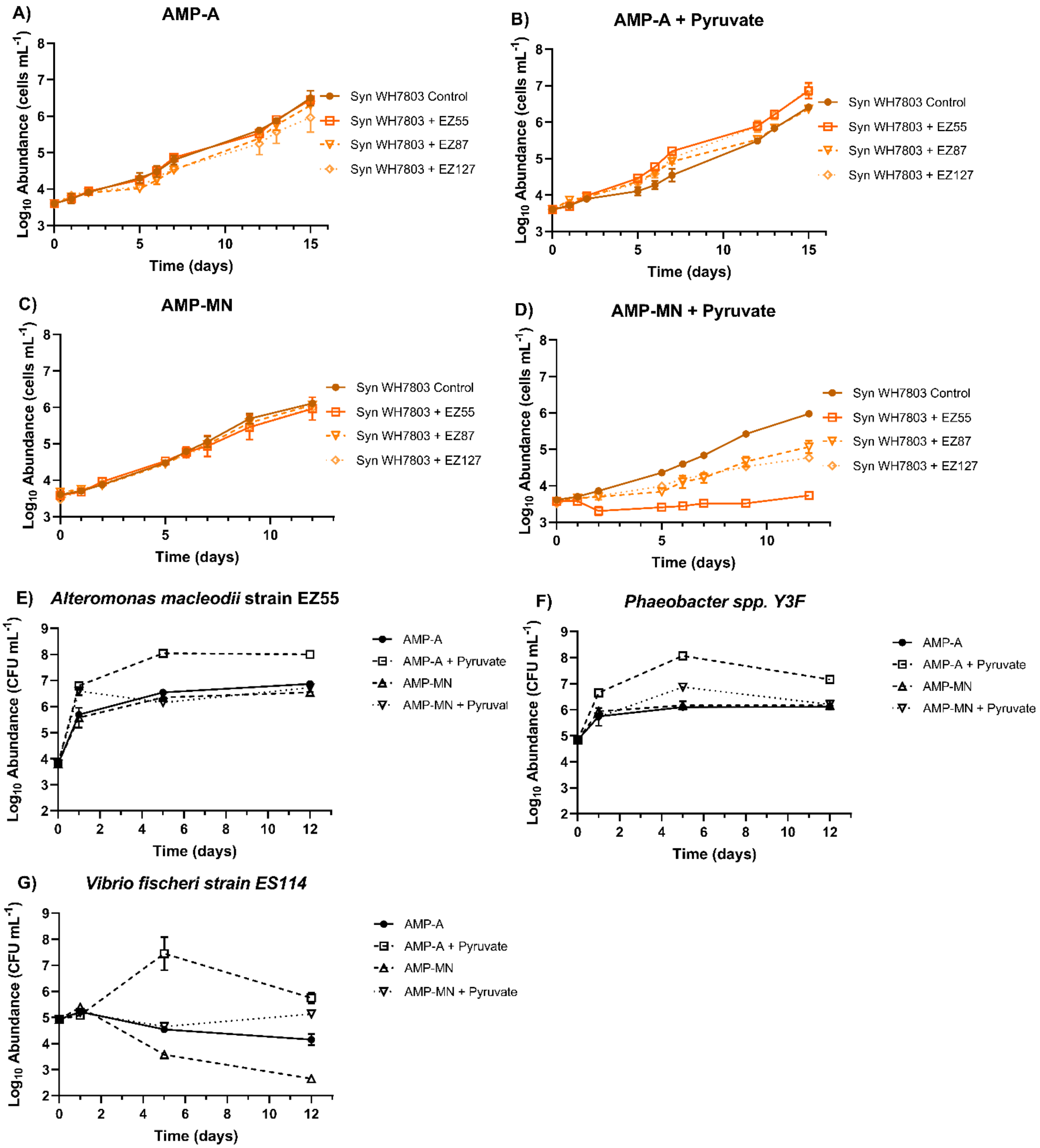
A)
AMP-A
Log10 Abundance (cells mL-1)
Time (days)
Syn WH7803 Control
Syn WH7803 + EZ55
Syn WH7803 + EZ87
Syn WH7803 + EZ127
B)
AMP-A + Pyruvate
Log10 Abundance (cells mL-1)
Time (days)
Syn WH7803 Control
Syn WH7803 + EZ55
Syn WH7803 + EZ87
Syn WH7803 + EZ127
C)
AMP-MN
Log10 Abundance (cells mL-1)
Time (days)
Syn WH7803 Control
Syn WH7803 + EZ55
Syn WH7803 + EZ87
Syn WH7803 + EZ127
D)
AMP-MN + Pyruvate
Log10 Abundance (cells mL-1)
Time (days)
Syn WH7803 Control
Syn WH7803 + EZ55
Syn WH7803 + EZ87
Syn WH7803 + EZ127
E)
Alteromonas macleodii strain EZ55
Log10 Abundance (CFU mL-1)
Time (days)
AMP-A
AMP-A + Pyruvate
AMP-MN
AMP-MN + Pyruval
F)
Phaeobacter spp. Y3F
Log10 Abundance (CFU mL-1)
Time (days)
AMP-A
AMP-A + Pyruvate
AMP-MN
AMP-MN + Pyruvate
G)
Vibrio fischeri strain ES114
Log10 Abundance (CFU mL-1)
Time (days)
AMP-A
AMP-A + Pyruvate
AMP-MN
AMP-MN + Pyruvate

**Fig. S4** – Growth of *Synechococcus* strain WH7803 in AMP-A and AMP-MN artificial seawater media with and without 500 µM sodium pyruvate in monoculture and coculture with individual heterotrophs (A-D): *Alteromonas macleodii* strain EZ55, *Vibrio fischeri* strain ES114 (EZ87), and *Phaeobacter* sp. strain Y3F (EZ127). Heterotroph abundances in all coculture treatments are shown (E-G). Error bars represent one standard deviation of the geometric mean (n=3).
